# The mouse claustrum synaptically connects cortical network motifs

**DOI:** 10.1101/2022.03.31.486634

**Authors:** Houman Qadir, Brent W. Stewart, Jonathan W. VanRyzin, Qiong Wu, Shuo Chen, David A. Seminowicz, Brian N. Mathur

## Abstract

Spatially distant areas of cerebral cortex coordinate their activity into networks that are integral to cognitive processing. A common structural motif of cortical networks is co-activated frontal and posterior cortical regions. Knowledge of the neural circuit mechanisms underlying such widespread inter-areal cortical coordination is lacking. Using anesthetized mouse functional magnetic resonance imaging (fMRI) we discovered that mouse frontal cortical functional connectivity reflects the common cortical network motif in its functional connectivity to posterior cortices, but also demonstrates significant functional connectivity with the claustrum. Exploring whether the claustrum may synaptically support such network architecture, we used a channelrhodopsin-assisted electrophysiological circuit mapping approach to assess the strength of synaptic connectivity of 35 unique frontal cortico-claustral-cortical connections through 1,050 subtype-identified claustrum projection neurons. We observed significant trans-claustral synaptic connectivity from the anterior cingulate cortex and prelimbic prefrontal cortex back to originating frontal cortical regions as well as to posteriorly-lying visual and parietal association cortices contralaterally. The infralimbic prefrontal cortex possessed significant trans-claustral synaptic connectivity with the posteriorly-lying retrosplenial cortex, but to a far lesser degree with visual and parietal association cortices. These data reveal discrete extended cortical pathways through the claustrum that are positioned to support cortical network motifs central to cognitive control functions.

## Introduction

The transfer of executive cortical information through subcortical structures that lead back to cortex is essential for cognition and the implementation of complex behavioral strategies (Rafal and Posner et al., 1987; Kraft et al., 2015; Voytek and Knight et al., 2010; Thompson et al., 1987, Packard and Knowlton 2002; Houk et al., 2007; Seger 2006). Such classical extended cortical systems include cortico-basal ganglia-cortical and cortico-thalamo-cortical loops. Delineating the specific directionality of flow of information through these multi-synaptic pathways has proven critical to advancing our understanding of their functional attributes (Sherman and Guillery 2002; Sherman 2017; Albin 1989; Bostan 2018; Aoki et al., 2019).

An understudied, yet significant, projection system emanating largely from frontal cortices also routes to the subcortical nucleus the claustrum. The claustrum (White and Mu et al., 2020; Atlan et al., 2018) and its frontal cortical input (White et al., 2018a) are required for optimal performance during cognitively demanding tasks. In humans, the claustrum is activated during execution of difficult, but not easy, versions of the multi-source interference attention task, which occurs coincidently with the emergence of task-positive cortical networks such as the fronto-parietal network (FPN) (Krimmel et al., 2019b). The FPN, along with the default mode network of task-negative cortical areas, are also functionally connected with the claustrum at rest (Krimmel et al., 2019b; Barrett et al., 2020). Given that cortical networks are initiated by frontal cortical regions (Grent-‘t-Jong and Woldorff et al., 2007), and cognitive control processes originate in frontal cortices (Botvinick, 2001; Shenhav, 2013; Miller and Buschman et al., 2007), the claustrum is positioned as a subcortical structure that may support cortical networks through discrete cortico-claustro-cortical pathways.

While evidence exists supporting claustrum functional connectivity with the salience network in rat (Smith, 2019) and a degree of synaptic connectivity to support this (Chia et al., 2020), further investigation of how the claustrum may provide a circuit mechanism supporting cortical network motifs composed of frontal and posterior cortical regions, such as task-positive and task-negative networks, is lacking. Characterizing a circuit mechanism supporting network communication may provide critical insight into myriad neuropsychiatric disorders in which the loss of network integrity predicts cognitive dysfunction, including addiction (Costumero et al., 2018), depression (Sylvester et al., 2013), and schizophrenia (Cole et al., 2011; Sheffield et al., 2015).

We analyzed the resting state functional connectivity (rsFC) of five frontal cortical seed regions of interest using mouse functional magnetic resonance imaging (fMRI) data to assess for claustrum functional connectivity. Testing the possible structural and synaptic connectivity underlying the functional connectivity observed using this approach, we examined 35 unique frontal cortico-claustral-cortical circuits using synaptic circuit mapping across 1,050 claustrum projection neurons of two physiologically distinct subtypes (White et al., 2018b). These data reveal distinct, primary information pathways through the claustrum reflecting a motif common in cortical networks underlying cognition.

## Results

### Mouse fMRI reveals rsFC between frontal cortical regions and claustrum

Both task-positive and task-negative networks are composed of specific frontal and posterior cortical regions. For example, the task-positive fronto-parietal network is composed of the cingulate cortex, dorsolateral prefrontal cortex, and posterior parietal cortex in humans (Sturm and Willmes et al., 2001; Chadick et al., 2011; Ptak, 2011; Hugdahl et al., 2015). The task-negative default mode network includes the ventromedial prefrontal cortex and posterior cingulate cortex (Raichle et al., 2001; Uddin et al., 2009). Previous imaging data reveal that such anti-correlated networks are conserved, to an extent, across rodents (Whitesell et al., 2021; Lu et al., 2012). Since cortical networks are initiated by their frontal cortical components (Grent-‘t-Jong and Woldorff et al., 2007), we sought to examine the functional connectivity of a host of well-characterized frontal regions in mice including anterior cingulate cortex (ACC), prelimbic prefrontal cortex (plPFC), infralimbic prefrontal cortex (ilPFC), the orbitofrontal cortex (OFC) and anterior insular cortex (aINS). Assessing whether these cortical areas possess functional connectivity with the claustrum we used a publicly available fMRI dataset (https://public.data.donders.ru.nl/dcmn/DSC_4180000.18_502_v1) acquired at 9.4 T (n = 51 mice). Following selection of unilateral (left) ROIs for all five frontal seed regions (Figure 1A), resting state functional connectivity (rsFC) maps for each frontal cortex seed exhibited significantly connected voxels surviving a conservative voxel-wise correction for multiple comparisons (FWE p < 0.05) within bilateral claustrum (CL) regions (Figure 1B-G). In addition, significantly connected voxels were also observed in posterior cortical regions, including the retrosplenial cortex (RSC), parietal association cortex (PtA), and visual cortex (V1/V2) (Figure 1H).

**Figure 1.**
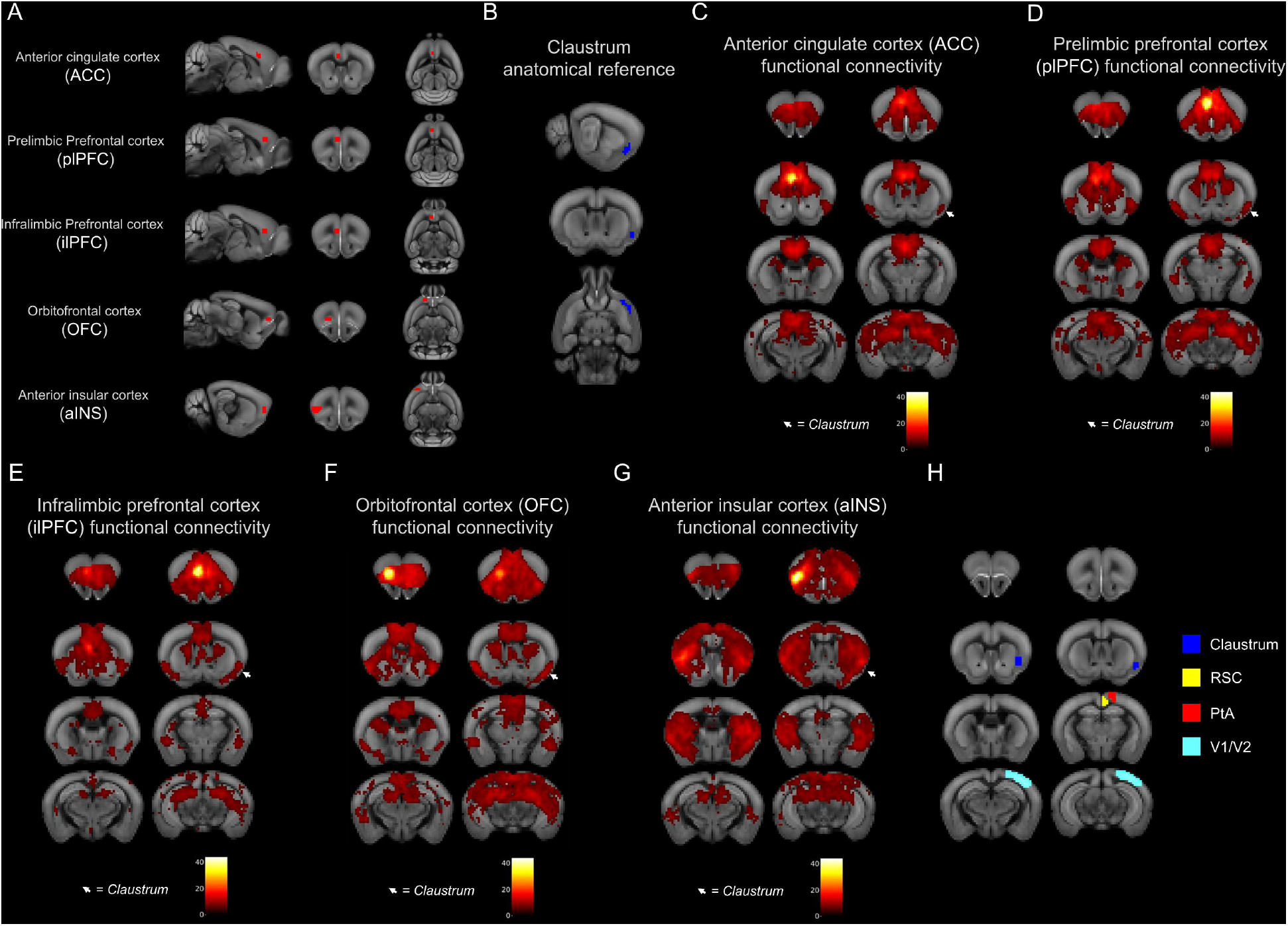
Frontal cortex seeds exhibit resting state functional connectivity (rsFC) with the claustrum. A) Sagittal (left), coronal (middle), and axial (right) views of unilateral anterior cingulate cortex (ACC), prelimbic prefrontal cortex (plPFC), infralimbic prefrontal cortex (ilPFC) orbitofrontal cortex (OFC), and anterior insular cortex (aINS) seed ROIs as drawn overlaid on the Allen Institute for Brain Science (AIBS) mouse template. B) Coronal slices displaying the rostro-caudal extent of the contralateral claustrum ROI overlaid on the AIBS mouse template. Slices correspond to those over which the rsFC heat maps of C) ACC, D) plPFC, E) ilPFC, F) OFC, and G) aINS are displayed (multiple comparisons-corrected voxel-wise FWE p < 0.05). Color bars indicate t-statistic values. White arrows indicate regions of rsFC overlap with contralateral claustrum. H) Coronal slices displaying claustrum with parietal cortices including retrosplenial cortex (RSC), parietal association cortex (PtA), and visual cortex (V1/V2).

### Claustrum projection neuron subtypes differ by firing properties and morphology

Functional connectivity analyses reveal voxels with timeseries significantly correlated with the seed region, and functional connectivity often reflects anatomical features (Greicius et al., 2009; Gordon et al., 2017). However, functional and anatomical connectivity do not necessarily correspond nor can be definitively interpreted as evidence of a direct influence of one brain region on another (Friston, 2011). Consequently, the rsFC data, while suggestive, do not allow conclusions regarding underlying synaptic connections nor cellular subtype specificity. We therefore investigated the strength of synaptic connectivity in distinct cortico-claustro-cortical circuits originating in the frontal cortex seed regions that target ipsilaterally back to the five frontal cortical regions as well as to the posterior cortical network regions identified in our rsFC data maps (Figure 1H).

Previous work suggested the existence of two potential projection neuron subtypes in the claustrum (White et al., 2018b). To confirm this, we sought to distinguish these neurons by both electrophysiological and morphological data. To do this, mice of both sexes received bilateral injections of an anterograde GFP-expressing virus (AAV5-hSyn-eGFP) in the ACC to outline the anatomical boundary of the claustrum of both hemispheres (White et al., 2018b). We recorded from claustrum projection neurons within the GFP-marked claustrum boundaries using a biocytin-filled internal recording solution to create three-dimensional reconstructions of the recorded neurons (Figure 2A). The identification of each claustrum projection neuron was determined based on burst-firing properties (Figure 2B): “type II” claustrum projection neurons burst fire following a brief 2ms depolarizing voltage step whereas “type I” neurons do not (White et al., 2018b). Following Sholl analysis of the reconstructed claustrum neurons, type II neurons exhibited a significantly greater dendritic length and number of dendritic intersections than type I neurons (Figure 2C). The increased number of intersections appeared in higher branch order numbers (Figure 2D). While type II gross dendritic morphology was more complex than that of type I neurons, type I neurons exhibited greater dendritic spine density compared to type II neurons (Figure 2E). Since both projection neuron subtypes significantly differed on physiological and morphological grounds, we heretofore tested each neuronal subtype for differences in cortico-claustro-cortical connectivity.

**Figure 2.**
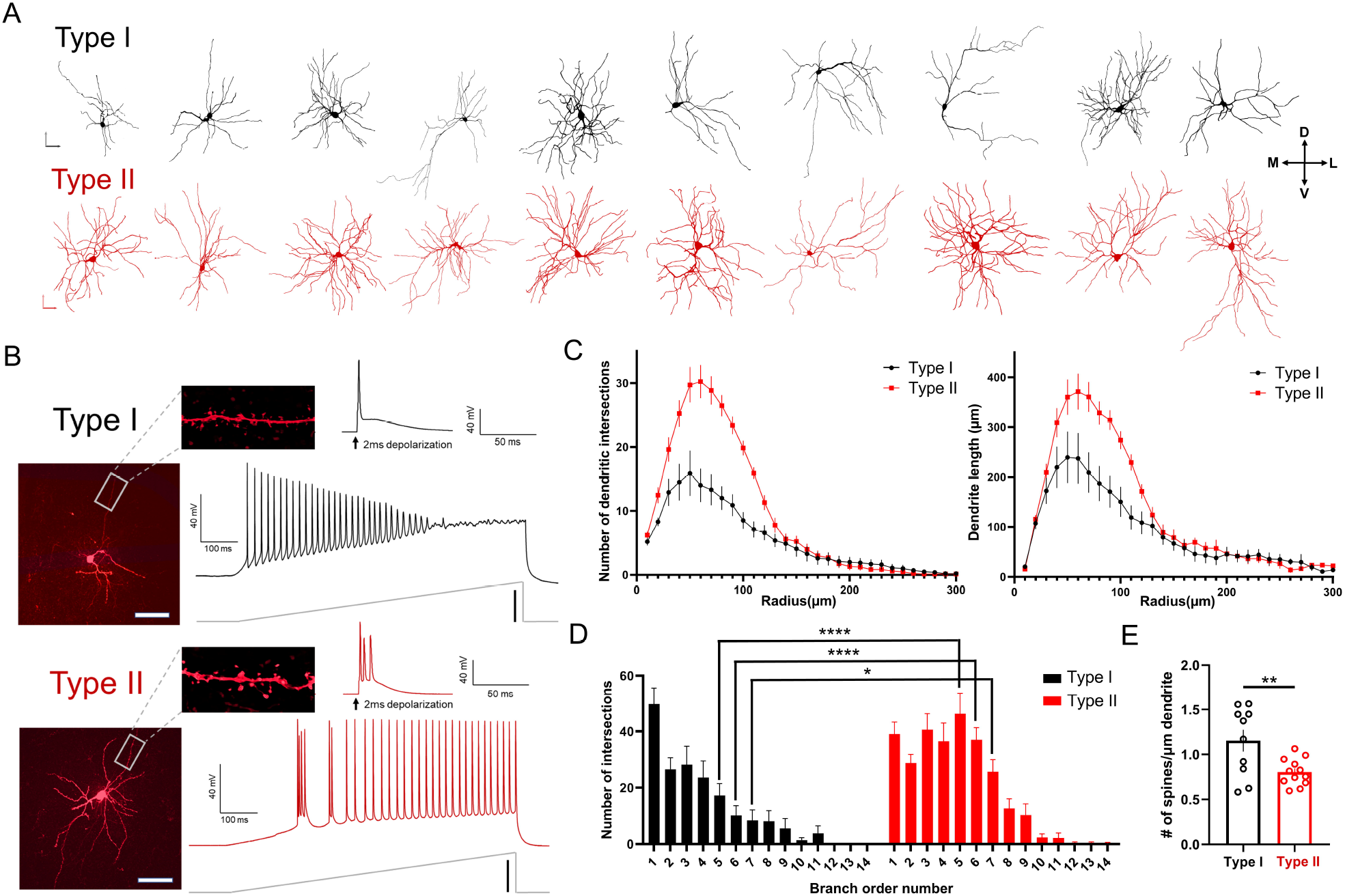
Type II spiny claustrum projection neurons dendritic morphology is more complex than Type I neurons. A) Top: Representative cell fill 3D reconstructions of Type I claustrum projection neurons. Bottom: Representative cell fill 3D reconstructions of Type II claustrum projection neurons (n=10 cells shown for each subtype). B) Top: Type I claustrum projection neuron delineated by lack of burst-firing following 2ms depolarization voltage step. Representative voltage trace following current-injection ramp. Bottom: Type II claustrum projection neuron delineated by presence of burst-firing following 2ms depolarization voltage step. Representative voltage trace following current-injection ramp. C) Type II claustrum neurons have increased number of dendritic intersections and increased dendrite length (Kruskal-Wallis test: P<0.0001; Type I: n=11 cells; Type II: n=12 cells). D) Type II neurons have increased number of intersections of higher branch order numbers (two-way ANOVA: F(13,308) = 3.97, P<0.0001, Bonferroni post hoc: P<0.01). E) Type I neurons have increased number of spines per μm of dendrite (unpaired-t test, P=0.008). Vertical scale bars: A) 100μm B) current ramp: 200pA. Horizontal scale bars: A) 100μm B) 200μm.

### Structural connectivity suggest multiple frontal cortico-claustral-cortical circuits exist

We next endeavored to test 35 possible frontal cortico-claustro-cortical circuits through both claustrum projection neuron subtypes based on our five input regions and seven output regions (Figure 3A). We bilaterally injected an anterograde eYFP virus (AAV5-hSyn-ChR2-eYFP) into various frontal cortical regions (Figure 3B), including the ACC (Figure S1A), plPFC (Figure S1B), ilPFC (Figure S1C), OFC (Figure S1D), and aINS (Figure S1E). A retrograde tdTomato virus (AAVrg-CAG-tdTomato) was also injected bilaterally in either the ACC (Figure S1F), plPFC (Figure S1G), ilPFC (Figure S1H), OFC (Figure S1I), aINS (Figure S1J), PtA (Figure S1K), V1/V2 (Figure S1L), and RSC (Figure S1M) to observe overlap between anterograde and retrograde labeling within the claustrum. We observed dense terminal anterograde expression throughout the rostral-caudal axis of the claustrum from the ACC (Figure S1A), plPFC (Figure S1B), and ilPFC (Figure S1C). Moderate eYFP expression was observed in the claustrum following bilateral injections in the OFC (Figure S1D) and sparse labeling was observed from the aINS in the claustrum (Figure S1E). Inputs arising from parietal sensory regions were not tested since optical stimulation from sensory cortices fails to elicit significant neuronal depolarization in claustrum projection neurons (White et al., 2018a). We found the fluorescent retrogradely labelled cell bodies in the claustrum targeted the ACC (Figure S1F), plPFC (Figure S1G), ilPFC (Figure S1H), OFC (Figure S1I), PtA (Figure S1K), V1/V2 (Figure S1L), and RSC (Figure S1M). We found no cells labelled in the claustrum following retrograde viral injections in the aINS (Figure 2I), which confirms the claustrum weakly projects to this area (Qadir et al., 2018).

**Figure 3.**
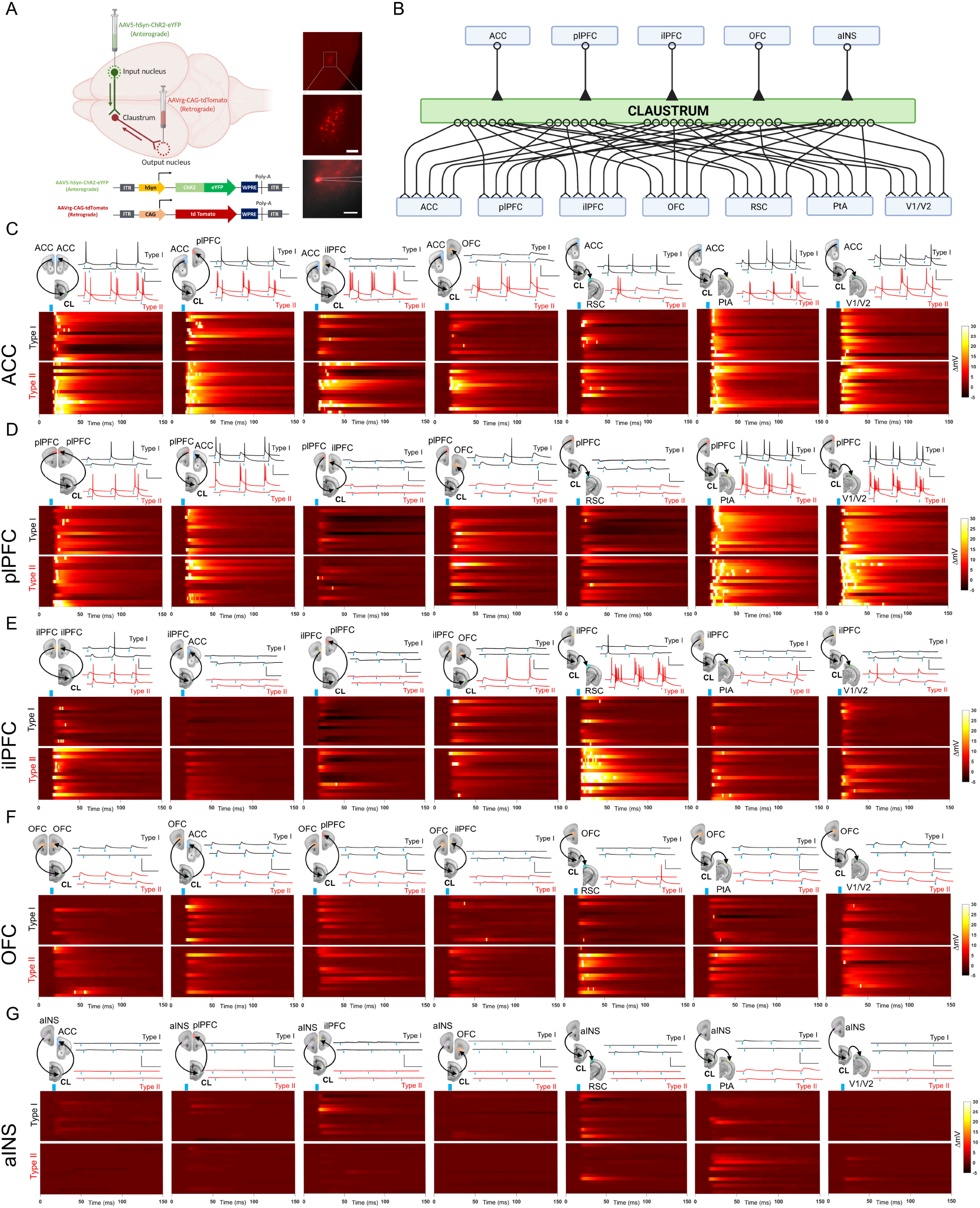
Functional channelrhodopsin-assisted circuit mapping of frontoclaustro-cortical circuits. A) Left: illustration of viral set up for structural circuit mapping method. AAV5-hSyn-ChR2-eYFP injected into the input cortical nucleus for anterograde terminal labeling and AAVrg-CAG-tdTomato injected in output cortical nucleus for retrograde soma labeling in claustrum. Right: representative image of fluorescently labeled spiny claustrum projection neurons for fluorescence-guided slice recordings. B) Cartoon diagram of all 35 fronto-claustro-cortical circuits tested with 5 input frontal cortical regions and 7 output regions. C) Top: Diagram of ex vivo ACC trans-claustral circuits tested (projecting to ACC, plPFC, ilPFC, OFC, RSC, PtA, and V1/V2 cortices respectively) and corresponding representative voltage traces for recorded type I and II claustrum neurons. Bottom: Heatmaps depicting average change in membrane potential across each recording trace following each light pulse stimulation (blue marker) for type I and II claustrum neurons. Only recordings with maximum light intensity (3mW) are shown. Circuits tested: ACC > CL > ACC; ACC > CL > plPFC; ACC > CL > ilPFC; ACC > CL > OFC; ACC > CL > RSC; ACC > CL > PtA; ACC > CL > V1/V2 (n=15 type I cells; n=15 type II cells each circuit). D) Diagram of ex vivo plPFC trans-claustral circuits tested (projecting to plPFC, ACC, ilPFC, OFC, RSC, PtA, and V1/V2 cortices respectively) and corresponding representative voltage traces for recorded type I and II claustrum neurons. Circuits tested: plPFC > CL > plPFC; plPFC> CL > ACC; plPFC > CL > ilPFC; plPFC > CL > OFC; plPFC > CL > RSC; plPFC > CL > PtA; plPFC > CL > V1/V2 (n=15 type I cells; n=15 type II cells each circuit). E) Diagram of ex vivo ilPFC trans-claustral circuits tested (projecting to ilPFC, ACC, plPFC, OFC, RSC, PtA, and V1/V2 cortices respectively) and corresponding representative voltage traces for recorded type I and II claustrum neurons. Circuits tested: ilPFC > CL > ilPFC; ilPFC> CL > ACC; ilPFC > CL > plPFC; ilPFC > CL > OFC; ilPFC > CL > RSC; ilPFC > CL > PtA; ilPFC > CL > V1/V2 (n=15 type I cells; n=15 type II cells each circuit). F) Diagram of ex vivo OFC trans-claustral circuits tested (projecting to OFC, ACC, plPFC, ilPFC, RSC, PtA, and V1/V2 cortices respectively) and corresponding representative voltage traces for recorded type I and II claustrum neurons. Circuits tested: OFC > CL > OFC; OFC> CL > ACC; OFC > CL > plPFC; OFC > CL > ilPFC; OFC > CL > RSC; OFC > CL > PtA; OFC > CL > V1/V2 (n=15 type I cells; n=15 type II cells each circuit). G) Diagram of ex vivo aINS trans-claustral circuits tested (projecting to ACC, plPFC, ilPFC, OFC, RSC, PtA, and V1/V2 cortices respectively) and corresponding representative voltage traces for recorded type I and II claustrum neurons. Circuits tested: OFC > CL > OFC; OFC> CL > ACC; OFC > CL > plPFC; OFC > CL > ilPFC; OFC > CL > RSC; OFC > CL > PtA; OFC > CL > V1/V2 (n=15 type I cells; n=15 type II cells each circuit). Horizontal scale bars: A) Top: 500μm Bottom: 200μm C-G) 100ms. Vertical scale bars: C-G): 40mV.

### Synaptic circuit mapping reveals distinct cortico-claustro-cortical circuits

To determine whether the structural connections observed indeed form synaptically-connected cortico-claustro-cortical pathways – and to what degree of strength they form - we used a channelrhodopsin-assisted long-range circuit mapping approach. We injected anterogradely transported AAV5-hSyn-ChR2-eYFP in each of the frontal cortical seed regions used previously (Figures 1 and 3): ACC, plPFC, ilPFC, OFC, and aINS. Retrogradely labeled tdTomato-positive claustrum projection neurons were recorded using whole-cell patch clamp for each cortico-claustro-cortical circuit (Figure 3A). Each neuron was first categorized as a type I or II claustrum neuron based on their action potential firing response to a brief depolarizing voltage step (Figure 2B).

Based on the area under the synaptic response curve (AUC) (Figure 4) and action potentials (APs) per light pulse synaptic strength metrics (Figure 5), we discovered four distinct frontal cortico-claustro-cortical circuits that terminated back on the originating cortical area on the contralateral side (e.g., left ACC > CL > right ACC) we termed “homoloquial circuits”. These circuits include ACC > CL > ACC (AUC = type I: 980.57 ± 234.03 mV*msec, APs/light pulse = 0.20 ± 0.09; type II: 1888.64 ± 621.51 mV*msec, 0.55 ± 0.21) (Figure 3C,4A,5A); plPFC > CL > plPFC (type I: 829.25 ± 283.61 mV*msec, 0.10 ± 0.0; type II: 1397.84 ± 400.05 mV*ms, 0.30 ± 0.14) (Figure 3D,4A,5A); ilPFC > CL > ilPFC (type I: 418.33 ± 82.48 mV*msec, 0.06 ± 0.03; type II: 1521.39 ± 379.03 mV*msec, 0.32 ± 0.11) (Figure 3E,4A,5A); and OFC > CL > OFC (type I: 334.97 ± 56.41 mV*msec, 0.00 ± 0.00, type II: 576.29 ± 143.41 mV*msec, 0.02 ± 0.01 (Figure 3F,4A,5A). An aINS homoloquial circuit was not tested as we found no claustrum neurons projecting to the aINS (Figure S1E).

**Figure 4.**
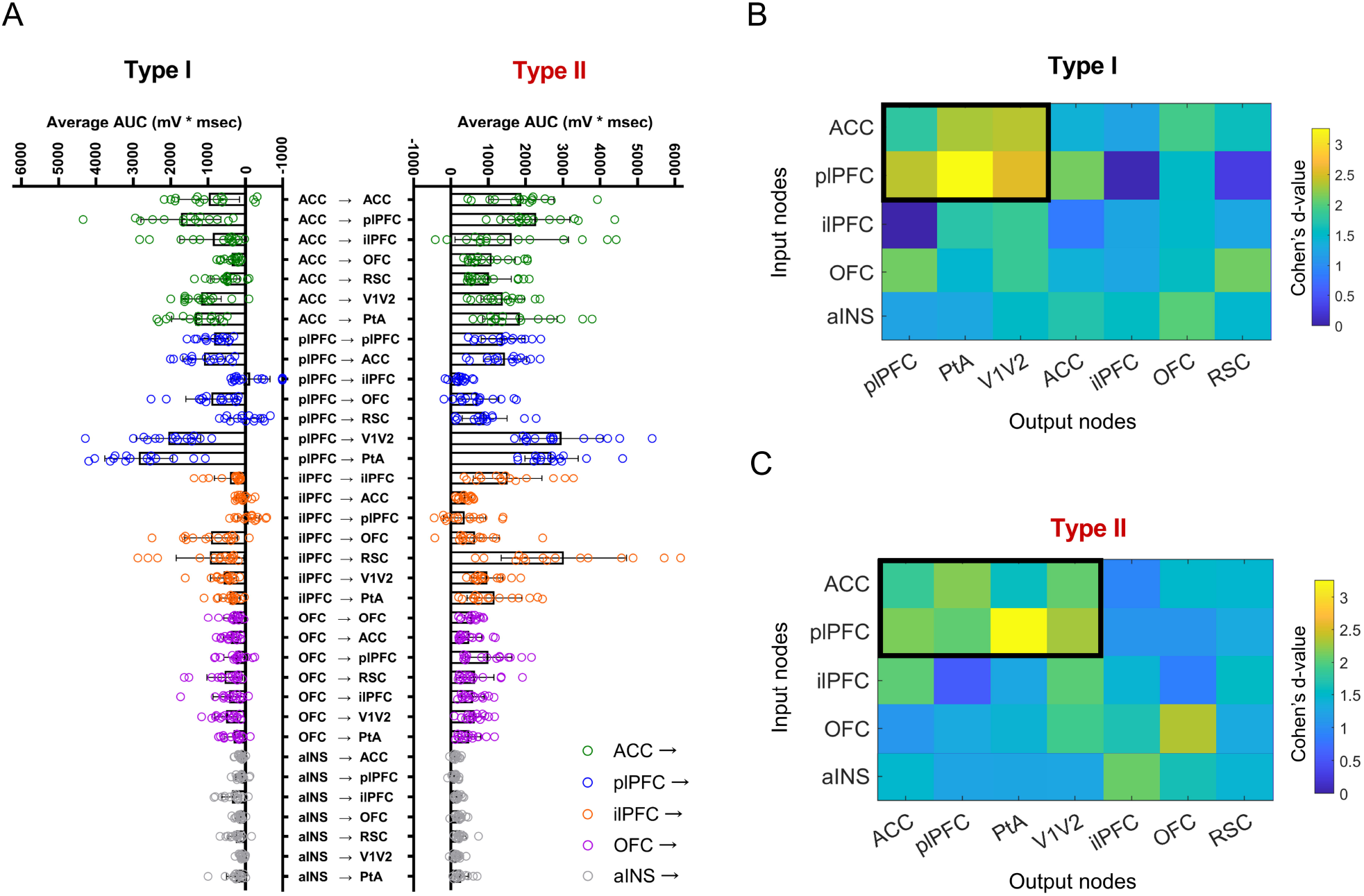
Average area under the curve metric shows specificity of circuit strength based on claustrum neuron output target region. A) Averaging the area under the curve of each voltage trace (for all light intensities) reveals differences in circuit strength across frontal trans-claustral circuits depending on type I and II claustrum projection neuron target (Kruskal Wallis test for multiple comparisons: P<0.0001; n=525 type I cells and n=525 type II cells total). B-C) Clustered strong frontal cortico-claustro-cortical circuits were detected by subgraph extraction (cluster marked in black) for type I and type II claustrum neuron subtypes. Type I: detected subgraph from ACC and plPFC to plPFC, PtA, and V1/V2 had high proportion of significant circuits. Type II: detected subgraph from ACC and plPFC to ACC, plPFC, PtA, V1/V2 had high proportion of significant circuits. Detected subgraphs P<0.001 under permutation test.

**Figure 5.**
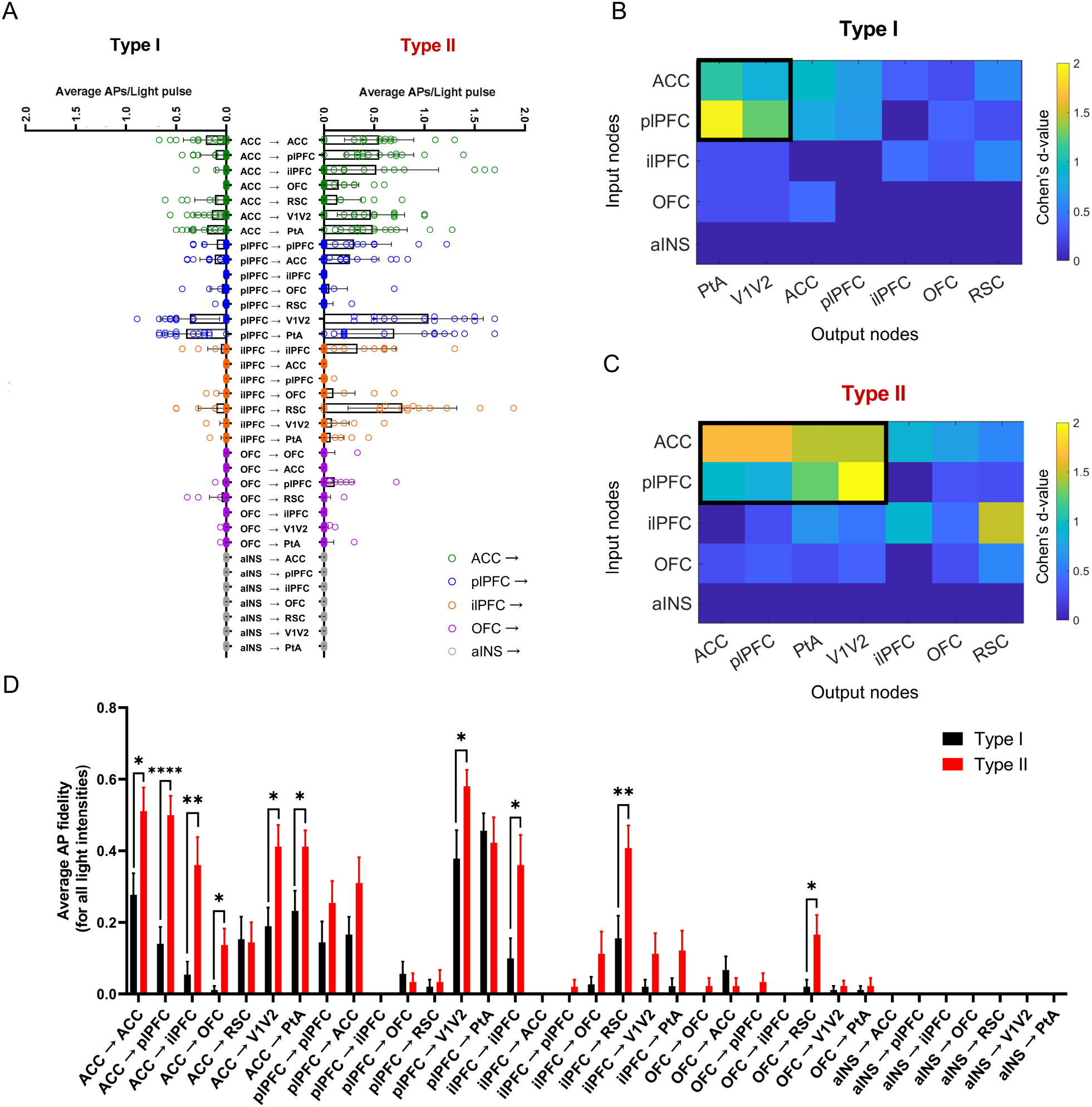
Average action potentials per light pulse metric shows specificity of circuit strength based on claustrum neuron output target region and firing fidelity is neuron subtype specific. A) Average action potentials per light pulse stimulation of each voltage trace (for all light intensities) reveals differences in circuit strength across frontal trans-claustral circuits depending on type I and II claustrum projection neuron target (Kruskal Wallis test for multiple comparisons: P<0.0001; n=570 type I cells and n=525 type II cells total). B-C) Clustered strong frontal cortico-claustro-cortical circuits were detected by subgraph extraction (cluster marked in black) for type I and type II claustrum neuron subtypes. Type I: detected subgraph from ACC and plPFC to PtA and V1/V2 had high proportion of significant circuits. Type II: detected subgraph from ACC and plPFC to ACC, plPFC, PtA, V1/V2 had high proportion of significant circuits. Detected subgraphs P<0.001 under permutation test. D) Select fronto-cortico-claustro-cortical circuits preferentially activate type II neurons compared to type I neurons: ACC > CL > ACC (Wilcoxon rank sum test: P=0.027); ACC > CL > plPFC (P<0.0001); ACC > CL > ilPFC (P=0.003); ACC > CL > OFC (P=0.006); ACC > CL > V1/V2 (P=0.016); ACC > CL > PtA (P=0.009); plPFC > CL > V1/V2 (P=0.041); ilPFC > CL > ilPFC (P=0.016); ilPFC > CL > V1/V2 (P=0.215); and OFC > CL > RSC (P=0.015). n=15 cells in each circuit for each subtype. Error bars: Standard Error.

The other 31 circuits tested were circuits where inputs originate in frontal cortices and synapse onto claustrum neurons that project ipsilaterally to a different cortical area from the originating frontal cortical region (termed “heteroloquial circuits”). ACC-originating heteroloquial circuits (Figure 3C,4A,5A) include: ACC > CL > plPFC (type I: 0.11 ± 0.06, 1726.36 ± 500.26 mV*msec; type II: 0.55 ± 0.23, 2283.24 ± 718.96 mV*msec); ACC > CL > ilPFC (type1: 0.03 ± 0.01, 874.50 ± 158.12 mV*msec; type II: 0.52 ± 0.18, 1630.08 ± 437.59 mV*msec); ACC > CL > OFC (type I: 0.00 ± 0.00, 365.18 ± 97.84 mV*msec; type II: 0.14 ± 0.09, 1092.71 ± 369.7 mV*msec); ACC > CL > RSC (type I: 0.11 ± 0.05, 543.90 ± 100.59 mV*msec; type II: 0.13 ± 0.07,1018.83 ± 228.81 mV*msec); ACC > CL > PtA (type I: 0.19 ± 0.10, 1605.65 ± 499.74 mV*msec; type II: 0.48 ± 0.22, 1846.68 ± 611.11 mV*msec); and ACC > CL > V1/V2 (type I: 0.14 ± 0.08, 1187.82 ± 339.33 mV*msec; type II: 0.46 ± 0.18, 1387.44 ± 427.60 mV*msec).

The plPFC-originating heteroloquial circuits (Figure 3D,4A,5A) include: plPFC > CL > ACC (type I: 0.11 ± 0.07, 1110.11 ± 329.41 mV*msec; type II: 0.26 ± 0.12, 1448.17 ± 422.10 mV*msec); plPFC > CL > ilPFC (type I: 0.00 ± 0.00, -126.12 ± 75.50 mV*msec; type II: 0.00 ± 0.00, 257.79 ± 33.61 mV*msec); plPFC > CL > OFC (type I: 0.04 ± 0.02, 910.73 ± 253.03 mV*msec; type II: 0.05 ± 0.03, 711.19 ± 171.64 mV*msec); plPFC > CL > RSC (type I: 0.00 ± 0.00, -5.03 ± 37.58 mV*msec; type II: 0.02 ± 0.01, 904.83 ± 172.07 mV*msec); plPFC > CL > PtA (type I: 0.40 ± 0.15, 2857.86 ± 1006.19 mV*msec; type II: 0.70 ± 0.26, 2698.31.15 ± 997.09 mV*msec); and plPFC > CL > V1/V2 (type I: 0.36 ± 0.12, 2063.42 ± 642.18 mV*msec; type II: 1.04 ± 0.36, 2963.15 ± 972.68 mV*msec).

The ilPFC-originating heteroloquial circuits (Figure 3E,4A,5A) include: ilPFC > CL > ACC (type I: 0.00 ± 0.00, 90.03 ± 10.76 mV*msec, type II: 0.00 ± 0.00, 393.51 ± 55.72); ilPFC > CL > plPFC (type I: 0.00 ± 0.00, -87.94 ± 34.25 mV*msec; type II: 0.01 ± 0.00, 367.92 ± 66.75 mV*msec); ilPFC > CL > OFC (type I: 0.02 ± 0.01, 921.15 ± 214.65 mV*msec; type II: 0.09 ± 0.03, 646.72 ± 141.09 mV*msec); ilPFC > CL > RSC (type I: 0.10 ± 0.04, 948.56 ± 203.32 mV*msec; type II: 0.78 ± 0.38, 3024.88 ± 900.08 mV*msec); ilPFC > CL > PtA (type I: 0.01 ± 0.01, 425.83 ± 69.29 mV*msec; type II: 0.09 ± 0.04, 1208.80 ± 261.53 mV*msec); and ilPFC > CL > V1/V2 (type I: 0.01 ± 0.01, 585.93 ± 95.61 mV*msec; type II: 0.07 ± 0.03, 974.24 ± 205.79 mV*msec).

The OFC-originating heteroloquial circuits (Figure 3F,4A,5A) include: OFC > CL > ACC (type I: 0.00 ± 0.00, 558.56 ± 120.27 mV*msec; type II: 0.01 ± 0.01, 645.47 ± 158.26 mV*msec); OFC > CL > plPFC (type I: 0.00 ± 0.00, 391.55 ± 72.17 mV*msec; type II: 0.01 ± 0.01, 496.05 ± 103.38 mV*msec); OFC > CL > ilPFC (type I: 0.00 ± 0.00, 437.82 ± 79.15 mV*msec; 0.00 ± 0.00, 590.06 ± 118.20 mV*msec); OFC > CL > RSC (type I: 0.01 ± 0.00, 260.83 ± 40.83 mV*msec, type II: 0.11 ± 0.06, 1006.56 ± 288.62 mV*msec); OFC > CL > PtA (type I: 0.00 ± 0.00, 318.96 ± 51.90 mV*msec; type II: 0.02 ± 0.01, 495.16 ± 158.10); and OFC > CL > V1/V2 (type I: 0.00 ± 0.00, 514.24 ± 135.05 mV*msec; type II: 0.01 ± 0.01, 634.90 ± 218.70 mV*msec).

Lastly, the aINS-originating heteroloquial circuits (Figure 3G,4A,5A) include: aINS > CL > ACC (type I: 0.00 ± 0.00, 133.14 ± 24.28 mV*msec; type II: 0.00 ± 0.00, 137.10 ± 14.48 mV*msec); aINS > CL > plPFC (type I: 0.00 ± 0.00, 126.87 ± 25.88 mV*msec; type II: 0.00 ± 0.00, 114.51 ± 23.34 mV*msec); aINS > CL > ilPFC (type I: 0.00 ± 0.00, 361.20 ± 66.63 mV*msec; type II: 0.00 ± 0.00, 196.16 ± 21.01); aINS > CL > OFC (type I: 0.00 ± 0.00, 170.44 ± 20.30 mV*msec; type II: 0.00 ± 0.00, 222.86 ± 21.19); aINS > CL > RSC (type I: 0.00 ± 0.00, 259.51 ± 46.23 mV*msec; type II: 0.00 ± 0.00, 257.16 ± 44.34 mV*msec); aINS > CL > PtA (type I: 0.00 ± 0.00, 261.55 ± 54.64 mV*msec; type II: 0.00 ± 0.00, 273.46 ± 93.64 mV*msec); and aINS > CL > V1/V2 (type I: 0.00 ± 0.00, 97.69 ± 21.15 mV*msec; type II: 0.00 ± 0.00, 150.73 ± 18.56 mV*msec). All seven aINS-originating circuits contained neurons that failed to elicit observable postsynaptic responses in the claustrum.

### Cortico-claustral synaptic strength depends on claustrum neuron subtype and postsynaptic target

We next applied a subgraph extraction-based cluster analysis to determine whether any group of circuits in this cortico-claustro-cortical circuit electrophysiological dataset emerge as statistically grouped to be physiologically relevant. Applying a permutation test with a significant p-value set at ≤ 0.001, we detected a subgraph of circuits that originate in the ACC and plPFC for both type I (Fig. 4B, 5B) and type II (Fig. 4C, 5C) neurons. Based on the average AUC metric the statistically significant clustered type I circuits include: ACC > CL > plPFC [Cohen’s D value = 1.66]; ACC > CL > PtA [2.23]; ACC > CL > V1/V2 [2.28]; plPFC > CL > plPFC [2.29]; plPFC > CL > PtA [3.25]; and plPFC > CL > V1/V2 [2.48]. AUC clustered type II circuits include: ACC > CL > ACC [2.24]; ACC > CL > plPFC [2.62]; ACC > CL > PtA [1.92]; ACC > CL > V1/V2 [2.49]; plPFC > CL > ACC [2.60]; plPFC > CL > plPFC [2.47]; plPFC > CL > PtA [3.91]; and plPFC > CL > V1/V2 [2.75]. Based on the average APs/light pulse metric the statistically significant clustered type I circuits include: ACC > CL > PtA [1.09]; ACC > CL > V1/V2 [0.85]; plPFC > CL > PtA [1.94]; and plPFC > CL > V1/V2 [1.27]. APs/light pulse clustered type II circuits include: ACC > CL > ACC [1.65]; ACC > CL > plPFC [1.66]; ACC > CL > PtA [1.46]; ACC > CL > V1/V2 [1.45]; plPFC > CL > ACC [0.93]; plPFC > CL > PtA [1.46]; and plPFC > CL > V1/V2 [2.00]. Notably, while ilPFC > CL > RSC (Fig. 4A,5A) exhibited connectivity, it did not reach statistical significance for clustering into ACC- and plPFC-originating circuits.

Based on the cluster analysis results, plPFC- and ACC-originating circuits, which both display cortico-claustro-cortical connectivity with V1/V2 and PtA, are statistically related. This contrasted with ilPFC-originating circuits that displayed weak connections to PtA and V1/V2 -projecting claustrum neurons (Figures 4 and 5). Instead, ilPFC connects significantly with RSC-projecting claustrum neurons, particularly through type II claustrum neurons. Extending this disparity plPFC activation induces hyperpolarizing response in claustrum neurons projecting to the ilPFC-favored target region the RSC and to claustrum projection neurons targeting ilPFC itself (Figure 4a). Further, ilPFC activation induces hyperpolarizing responses in plPFC-projecting claustrum neurons.

Lastly, we compared the postsynaptic responses of type I versus type II neurons for each of the 35 circuits tested (Figure 5D). To account for burst firing properties of type II neurons, we transformed the data into binary responses (0 = no action potentials and 1 = at least one action potential). Using this metric, we discovered a total of 10 circuits in which type II neurons significantly fired more than type I neurons: ACC > CL > ACC (type I: 0.28 ± 0.06, type II: 0.51 ± 0.07); ACC > CL > plPFC (type I: 0.14 ± 0.05, type II: 0.50 ± 0.05); ACC > CL > ilPFC (type I: 0.05 ± 0.04, type II: 0.36 ± 0.19), ACC > CL > PtA (type I: 0.23 ± 0.06, type II: 0.41 ± 0.05); ACC > CL > V1/V2 (type I: 0.19 ± 0.05, type II: 0.41 ± 0.06); ACC > CL > OFC (type I: 0.00 ± 0.00, type II: 0.13 ± 0.04); plPFC > CL > V1/V2 (type I: 0.38 ± 0.08, type II: 0.58 ± 0.05); ilPFC > CL > ilPFC (type I: 0.10 ± 0.06, type II: 0.36 ± 0.08); ilPFC > CL > RSC (type I: 0.16 ± 0.06, type II: 0.41 ± 0.06); and OFC > CL > RSC (type I: 0.02 ± 0.02, type II: 0.17 ± 0.05). Most circuits that preferentially drove AP firing in type II neurons over type I neurons were from ACC-originating circuits. No circuits tested preferentially activated type I neurons significantly more than type II neurons.

## Discussion

Mouse fMRI revealed frontal cortical region functional connectivity with claustrum and with posterior sensory and association cortices, suggesting underlying synaptic connectivity. Channelrhodopsin-assisted circuit mapping experiments uncovered two types of cortico-claustro-cortical circuits: homoloquial circuits that originate from frontal cortical regions and relay back to frontal regions and heteroloquial circuits that originate in frontal cortices and relay to posterior sensory regions. We found that the two physiologically distinct subtypes of claustrum projection neurons, which are also morphologically distinct, differentially support trans-claustral cortico-claustro-cortical circuits. However, frontal cortical inputs onto claustrum projections differ in strength depending on the output region of a given claustrum neuron regardless of subtype. For example, plPFC > CL > PtA, plPFC > CL > V1/V2, ACC > CL > plPFC, and ilPFC > CL > RSC circuits are significantly stronger than all other ACC-, plPFC-, and ilPFC-originating circuits as well as all OFC- and aINS-originating circuits tested. These data indicate that the claustrum is synaptically configured to allow for flow of information from frontal cortices back to frontal cortices, as well as to posterior cortices, in a circuit- and cell-type-specific manner that reflects whole-brain imaging functional connectivity data.

Like basal ganglia and thalamic structures, the present data describe an extended cortical system involving significant cortical input to a subcortical nucleus – the claustrum - that returns processed information back to cortex. The primary difference with the basal ganglia and thalamic nuclei, however, is that the claustrum provides input back not only to originating frontal cortical nuclei, but to geographically distant areas from frontal cortices, including parietal cortical structures. Higher-order thalamic structures such as the pulvinar, lateral posterior nuclei, and mediodorsal nuclei, receive converging input from layer 5/6 prefrontal and sensory cortical projection neurons (Collins et al., 2018; Groh et al., 2014), which in turn propagate incoming signals back to superficial layers 2/3 of the prefrontal cortex (Collins et al., 2018). This contrasts with the claustrum, which projects to layers 2/3, 5 and 6 in frontal cortices (Jackson et al., 2018; White et al., 2018b).

The common claustro-cortical input to both frontal and posterior cortices positions this structure to coordinate inter-areal cortical activity. Indeed, Narikiyo and colleagues (Narikiyo et al., 2020) showed that mouse claustrum activation synchronizes cortical activity. Thus, the major extended cortical communications network revealed herein, together with findings that the claustrum is functionally connected with human cortical networks (Krimmel et al., 2019b; Barrett et al., 2020), supports the notion that the claustrum may be a central subcortical support system for cortical network function. Moreover, psilocybin, an agonist of serotonin 2A receptors, which are highly expressed in claustrum (Pazos et al., 1985), disrupts claustrum activity, cortical network integrity, and claustrum functional connectivity with cortical networks in human subjects (Barrett et al., 2020).

In addition to the significant cluster of ACC and plPFC projections that innervated claustrum neurons that, in turn, project to visual cortices and PtA, we found that the ilPFC > CL > RSC pathway through predominantly type II neurons was robust. This strongly contrasts with our plPFC > CL > RSC results where most RSC-projecting claustrum neurons hyperpolarized in response to plPFC afferent stimulation. This stark synaptic connectivity contrast between ilPFC- and plPFC-driving circuits, may suggest that the claustrum supports discrete network states. Considering that human task-positive and task-negative cortical networks are anti-correlated (Fox et al., 2005; Uddin et al., 2009; Riemer et al., 2020), and that ilPFC and RSC are identified as two major putative nodes of the default mode network in mouse (Stafford et al., 2014), whereas the plPFC and PtA are putative mouse homologs of major nodes in the human fronto-parietal network (Hwang, 2021; Laubach et al., 2018), the present data may support the idea of the claustrum acting as a relay system sculpting defined cortical networks.

Our data suggest that the claustrum differentially relays frontal cortical signals in a claustrum projection neuron subtype-dependent manner. For example, we found that inputs arising from the ACC preferentially activate type II neurons significantly more than type I neurons. Although further work is needed to define the functional significance of these pathways, we speculate that since synchronized rhythms are a major hallmark of networks (Buschman et al., 2012), an executive cortical structure may “jump-start” specific networks states through the burst-firing properties of type II claustrum neurons. This notion fits with recent data showing claustrum neuron ensembles not being modulated by “bottom up” sensory inputs, but rather, are synchronized to preferring contralateral licks in a sensory selection task (Chevée et al., 2022). Given the transient firing nature of claustrum projection neurons (White et al., 2018b) and the ability for single claustrum neurons to project to multiple functionally related brain regions (Marriott et al., 2020), the claustrum may function to switch, but not maintain, cortical network states upon cognitive demand. This is supported by the finding that significant claustrum activation is observed when task-negative networks diminish and task-positive networks emerge at the beginning of a complex cognitive task (Krimmel et al., 2019b).

The present findings do not rule out the existence of other cortico-claustro-cortical or sub-cortical-claustro-cortical pathways. However, considering that the majority of input to the claustrum is cortical, and of that the majority arises from the frontal cortices (Wang et al., 2018), the present results highlight what is likely the bulk of information flow through claustrum. While the connections defined here support cortical network motif architectures, they also suggest that frontal cortical areas may communicate with one another through the claustrum, perhaps for dynamic control of downstream network states. Taken together with the thalamic nuclei and cortico-cortical connections that support putative default mode network connectivity in mouse (Whitesell et al., 2021), the cortical source, claustrum cell type-, and cortical target-specific pathways defined here all likely cooperate to support cortical network states for optimal cognitive performance.

## Acknowledgments

This work was supported by National Institute on Alcohol Abuse and Alcoholism grants R01AA024845 (B.N.M.) and R01AA028070 (B.N.M.), and National Institute of Mental Health grant F31MH126465 (H.Q).

## Materials and methods

### Animals

5 C57BL/6J (wild type) mice of both sexes were used for neuron 3D reconstruction experiments. 40 wild-type mice were used for circuit histology experiments. 175 wild-type mice were used for all ex vivo channelrhodopsin circuit mapping whole-cell patch clamp experiments (5 mice per circuit). Mice used for all ex vivo experiments were 12-16 weeks of age and were group-housed with food and water available ad limitum and on a 12 hour day/night light cycle beginning at 07:00 and all patch-clamp experiments were performed during the light cycle. This study was performed in accordance with the National Institutes of Health Guide for Care and Use of Laboratory Animals and the University of Maryland, School of Medicine, Animal Care and Use Committee.

### Stereotaxic Procedures and Viral vectors

Mice were anesthetized via inhalation of 3.5% isoflurane and placed in a mouse stereotaxic frame while anesthesia was maintained with 1% isoflurane inhalation. A stereotaxic drill was used to drill small openings in the mouse skull above brain regions prior to viral injection. 250nl of an anterograde adeno-associated virus (AAV) vector expressing a green fluorescent protein under the hSyn (human synapsin) promoter (AAV5-hSyn-eGFP; Addgene) was injected into ACC to fluorescently mark the anatomical boundary of the claustrum (White et al., 2017, Qadir et al., 2018) in order to cell fill spiny claustrum projection neurons for 3D reconstruction analysis. Relative to bregma, the coordinates used for ACC injections were anterior-posterior (AP): +1.0mm, medial-lateral (ML): ± 0.3mm, dorsal-ventral (DV): -1.1mm. For all CRACM slice electrophysiology experiments, 200nL injections into the input nucleus were performed bilaterally using an AAV vector expressing channelrhodopsin (AAV5 hSyn-ChR2-eYFP; Addgene) and simultaneously injected 150nL of a retrograde AAV expressing a td Tomato tag under the CAG (chicken beta-actin) promoter (rgAAV-CAG-td tomato; Addgene) (Tervo et al., 2016) into the output nucleus to fluorescently label claustrum projection neurons projecting to the target region. Exactly 4 weeks of virus incubation was given before mice were euthanized and brain slices were taken for ev vivo cellular recordings. Coordinates for the following brain regions were used for CRACM experiments: ACC: (see above); plPFC: (AP = +2.0mm, ML = ±0.4mm, DV = - 1.2mm); ilPFC: (AP = +1.78, ML = ±0.3mm, DV = -2.2mm) OFC: (AP = +2.6mm, ML = ±1.1mm, DV = -1.8mm); aINS: (AP = +1.94mm, ML = ±2.5mm, DV = -3.5mm); PtA: (AP = -1.9mm, ML = ±1.4mm, DV = -0.4mm); V1/V2: (AP = -2.9mm, ML = ±2.05mm, DV = - 0.4mm); RSC: (AP = -1.6mm, ML = ±0.3mm, DV = -0.5mm). DV coordinates were measure from top of brain surface.

### Histology

Mice were overdosed on isoflurane gas and perfused with room temperature 0.1M phosphate-buffered solution (PBS), pH 7.3, and then with ice-cold 4% paraformaldehyde (PFA) solution in PBS, 10 days after viral injection surgery. After extraction, the brains were post-fixed in 4% PFA solution overnight. 50 μm thickness slices were obtained using the Integraslice 7550 MM vibrating microtome (Campden Instruments, Loughborough, England), and were stored at 4°C in 0.1M PBS. The slices were mounted onto 25 × 75 × 1 mm frosted microscope slides (Thermo-Scientific, Waltham, MA, United States) using 125 uL ProLong Gold antifade reagent (Invitrogen) as the mountant. The slides were imaged using a Nikon fluorescence microscope (Nikon, Minato, Tokyo, Japan) with images obtained using both 4X and 10X magnification objectives.

### Resting State Functional Connectivity fMRI analysis

Mouse resting state fMRI data was obtained from a publicly available dataset (https://public.data.donders.ru.nl/dcmn/DSC_4180000.18_502_v1, https://public.data.donders.ru.nl/dcmn/DSC_4180000.18_502_v1/LICENSE.txt). The specific data used consisted of scans from an experiment testing the effects of a model of psychosocial stress on male, wild-type, C57BL/6 mice aged 3 months (Grandjean et al., 2016). We therefore used only pre-intervention baseline scans from all 51 animals, both control and experimental, available online.

Full details on data acquisition are described in (Grandjean et al., 2016). Briefly, anesthesia was induced with 3.5% isoflurane, mice were ventilated at 80 breaths per minute, and anesthesia was maintained with a combination of pancuronium bromide, medetomidine, and a gradual reduction to 0.5% isoflurane. Scans were acquired with a Bruker 94/30 Biospec spectrometer (Bruker BioSpin MRI, Ettlingen, Germany) operating at 9.4 T. Resting state fMRI scans consisted of 6 minutes of blood oxygenation level-dependent (BOLD) gradient-echo echo planar images acquired using repetition time TR = 1000 ms, echo time TE = 9.2 ms, flip angle FA = 90°, matrix size MS = 90 × 70, field of view FOV = 20 × 17.5 mm^2^, slice number NS = 12, slice thickness ST = 0.5 mm, slice gap SG = 0.2 mm, and bandwidth BW = 250,000 Hz.

Full details on preprocessing are described in (Mandino et al., 2021). Briefly, anatomical images were registered to the Allen Institute for Brain Science (AIBS) mouse template (https://atlas.brain-map.org/). Functional images were despiked (3dDespike), motion-corrected (3dvolreg), corrected for B1 field, denoised, brain-masked, registered linearly to corresponding anatomical images, and band-pass filtered (3dBandpass, 0.01–0.25 Hz).

The five unilateral frontal cortical seed regions of interest (ROIs) and the contralateral claustrum were drawn using FSLeyes. For each ROI, the AIBS template used for preprocessing was loaded in the software, an empty 3D mask with the same dimensions was generated and overlaid on the template, and the voxels of the ROI were hand-selected using the anatomical knowledge of the authors, Paxinos and Franklin’s the Mouse Brain in Stereotaxic Coordinates, Compact (2008), and anatomical landmarks visible in the AIBS template. The seed and claustrum ROIs as drawn can be seen in Fig. 1a.

Analyses were performed in SPM12. To assess whole brain functional connectivity of the seed regions, the SPM toolbox MarsBar was used to extract each animal’s mean BOLD signal timeseries from each seed ROI, and individual General Linear Models were produced in SPM for each animal/ROI consisting of the mean ROI timeseries and 6 motion parameters as regressors.

To determine significant functional connectivity, one sample t-tests were performed on resting state second-level contrast maps masked with the AIBS template binary mask. To correct for multiple comparisons, we imposed an FWE-corrected voxel-wise significance threshold of p < 0.05. Data are publicly available, and code and ROI files are available upon request.

Following the synaptic connectivity experiments, 3 contralateral output ROIs – RSC, PtA, and visual cortex - were drawn in FSL using the same process used for the unilateral input and claustrum ROIs described above. SPM’s ImCalc function was used to generate separate images representing the overlap of one of these output ROIs or the claustrum ROI and the FWE p < 0.05 rsFC maps for ACC, plPFC, and ilPFC input regions. Images of input region rsFC overlap with claustrum and contralateral output regions in Fig. 6b-d were generated by overlaying the AIBS mouse brain template, an input region rsFC map, and corresponding claustrum and output region overlap images in FSL.

### 3D reconstruction

Type I and type II claustrum projection neurons were recorded under whole-cell current clamp conditions. Respective neurons were recorded with a potassium-based solution (290-295 mOsm; pH) with 5% concentration biocytin to allow for proper cell fill into the soma and distal dendrites. Immediately following electrophysiological recording, slices were fixed in 4% paraformaldehyde overnight at 4°C. The next day, slices were washed in 0.1 M phosphate buffered saline (PBS) 3 × 20 min and blocked with 1% bovine serum albumin (BSA) in PBS + 0.3% Triton X-100 (PBS-T) for 2 h at room temperature. Slices were incubated with Alexa Fluor 594-streptavidin (Invitrogen, #S32356) (1:1000, 1% BSA in 0.3% PBS-T) overnight at 4°C. The next day, slices were washed in PBS 3 × 20 min, mounted on slides, and coverslipped in ProLong Diamond Antifade (Invitrogen, #P36965).

Confocal images were acquired with a Nikon A1 microscope equipped with 488 and 561 lasers. For neuronal reconstructions, slices were first imaged for both GFP and Alexa Fluor 594 expression to verify the position of each neuron within the claustrum. Neurons were then imaged using a 40x (0.95 NA) objective with a lateral resolution of 0.310 μm per pixel and a 0.727 μm z-step. For dendritic spine reconstruction and quantification, two dendrites were imaged per cell using a 100x (1.46 NA) oil-immersion objective with a lateral resolution of 0.05 μm per pixel and a 0.10 μm z-step. Dendrites were randomly chosen, taking care to avoid broken or proximal (<50 μm from cell body) sections, and a roughly 50 μm section was imaged for analysis. Prior to spine analysis, raw images were denoised using the Nikon Elements Denoise.ai algorithm.

Semi-automated neuronal reconstruction was performed using Neurolucida 360 (version 2020.1.1) with directional kernels algorithm. Dendritic spine analysis was conducted using a semi-automated analysis method using Biplane Imaris (version 9.5.1). The Filaments module was used to first reconstruct sections of dendrite then detect spines. Spine detection was edited for accuracy when necessary by an experimenter blind to cell type.

### Ex vivo brain slice preparation for slice electrophysiology

Mice between the ages of 5-8 weeks were surgically injected with both viruses bilaterally and following four weeks of virus incubation were euthanized for ex vivo recordings. Following anesthetization, mice were decapitated, and the brains were extracted. 250 μm coronal sections were sliced using a Leica VT1200 vibrating microtome in a high-sucrose artificial cerebrospinal fluid (aCSF) solution. The high-sucrose cutting aCSF solution was kept ice cold, continuously bubbled with carbogen (95% O_2_, 5% CO_2_), and was comprised of 194 mM sucrose, 30 mM NaCL, 4.5 mM KCL, 1 mM MgCl_2_, 26 mM NaHCO_3_, 1.2 mM NaH_2_PO_4_, and 10 mM D-glucose. Sections were incubated after slicing for 30 min at 33 °C in carbogen-bubbled aCSF (315–320 mOsm) that contained 124 mM NaCl, 4.5 mM KCl, 2 mM CaCl_2_,1 mM MgCl_2_, 26 mM NaHCO_3_,1.2 mM NaH_2_PO_4_, and 10 mM D-glucose. Brain slices were incubated at room temperature until whole-cell patch-clamp recordings, and patch recordings were performed in the same aCSF formulation used for incubation.

### Whole-cell current clamp recordings

Whole-cell recordings were performed at 29–31 °C using borosilicate glass recording pipettes of 3–7 MΩ resistance. For recordings performed in a current clamp configuration, recording pipettes were filled with a potassium-based solution (290–295 mOsm; pH 7.3) composed of 126 mM potassium gluconate, 4 mM KCl, 10 mM HEPES, 4 mM ATP-Mg, 0.3 mM GTP-Na and 10 mM phosphocreatine. Clampex software (Version 10.4; Molecular Devices) was used for all electrophysiological recordings. Recordings were filtered at 2 kHz and digitized at 10 kHz using MultiClamp 700B software (Molecular Devices). Claustrum projection neuron type was determined via a 5 ms depolarization step while recording in current-clamp mode to determine burst firing properties (Type I: no burst fire; Type II: burst fire). Following this protocol, membrane capacitance values were also recorded to confirm the characterization of neuron type (Type I: ∼75-130 pF; Type II: ∼130-200 pF) (White et al., 2018). For all CRACM experiments, three 5ms 470nm blue light pulses with 150ms intervals were given to evoke presynaptic transmitter release while recording from fluorescently labeled claustrum projections (Petreanu et al., 2007).

### Circuit mapping data analysis and statistics

Three 5ms light pulses were delivered to optically stimulate ChR2-expressing glutamatergic afferents arising from a given cortical region to drive postsynaptic responses. To quantify the degree of the postsynaptic response in each circuit, we used: 1) average action potentials (APs) per light stimulation intensity and 2) area under the curve (AUC) for each postsynaptic recording trace. For AUC, a higher positive AUC value represents a postsynaptic depolarization, reflecting an excitatory postsynaptic potential (EPSP) or an AP. Conversely, a negative AUC value reflects a postsynaptic hyperpolarization. Notably, using the same potassium-based internal solution used throughout, we found that the reversal potential for inhibitory synaptic currents at both type I and II neuron synapses was -75mV (Figure S2A-C). Electrophysiology data were analyzed using Clampex software (version 11.0.3). Area under the curve values were converted from voltage values for each trace in Clampex data table files into excel files. Values for each circuit, cell, and light intensity were analyzed using MatLab (version 2019a) using trapezoidal integration for area under the curve to access activation/inactivation of each cell following presynaptic stimulation. Representative heatmaps were averaged using Microsoft excel and plotted in MatLab. Multiple comparison statistical analyses were performed using GraphPad Prism software (Prism 8). Subgraph extraction analyses was performed by determining the p-value of a subgraph through the probability to obtain the subgraph under the null hypothesis that none of the circuits and edges are significant (approximated by permutation test). Cohen’s d values for all the data points were calculated using MatLab.

## Figure legends

**Figure S1. Structural connectivity of putative frontal cortico-claustral-cortical loops**

A-E) Top panels: representative photomicrographs of anterograde eYFP virus injection sites in the A) ACC, B) plPFC, C) ilPFC, D) OFC, and E) aINS. Bottom panels: representative photomicrographs of anterograde terminal expression in the claustrum from respective frontal regions. Structural inputs were strongest from the ACC and plPFC with moderate inputs from the OFC. Very weak inputs were observed from the aINS based on anterograde viral expression in the claustrum. F-K) Top panels: representative photomicrographs of retrograde td-Tomato virus injection sites in the ACC (F), plPFC (G), OFC (H), aINS (I), parietal association cortex (PtA) (J), and visual cortex (V1/V2) (K). Bottom panels: representative photomicrographs of retrograde td-Tomato expression in the claustrum. Highest cell body density was found in ACC and plPFC cases followed by dense expression of claustrum cell projection neurons targeting the OFC, PtA and V1/V2 cortices. Very few to no claustrum neurons project to the aINS based on lack of retrograde labeling. (n=3 per circuit case). Horizontal scale bars: B-K) 500μm.

**Figure S2. Reversal potential of GABA is lower than the average resting membrane potential for both claustrum projection neuron subtypes**. A) Input/output curve across multiple membrane potentials reveals a reversal potential of GABARs to be approximately -75mV for type I and B) type II neurons. C) There is no significant difference between type I and II GABAR reversal potential. Horizontal scale bars: A-B) 20ms. Vertical scale bars: A-B) 200pA.

**Figure S3. Ex vivo circuit-mapping average area under the curve and number of action potentials per light pulse raw data values table**. A) Average area under the curve values (left) and actional potentials per light pulse values (right) for each light intensity tested (0-3mW) of all type I claustrum neurons (n=525 cells). B) Average area under the curve values (left) and actional potentials per light pulse values (right) for each light intensity tested (0-3mW) of all type II claustrum neurons (n=525 cells). Average action potentials per light stimulation values were used to determine circuit strength classifications based on action potential firing rates. Error shown is in standard error. Units: ** = mV x msec.

**Figure S4. P-value matrices for average area under the curve and action potentials per light pulse multiple comparisons test show specificity in claustrum neuron activation depending on projection output target region**. A-B) Kruskal Wallis test for multiple comparisons P-value matrix for average area under the curve for type I (A) and type II (B) trans-claustral circuits. C-D) Kruskal Wallis test for multiple comparisons P-value matrix for average action potentials per light pulse for type I (C) and type II (D) trans-claustral circuits.

