## Supplementary figures and images for "The mouse claustrum synaptically connects cortical network motifs"

### Supplemental Figure S1

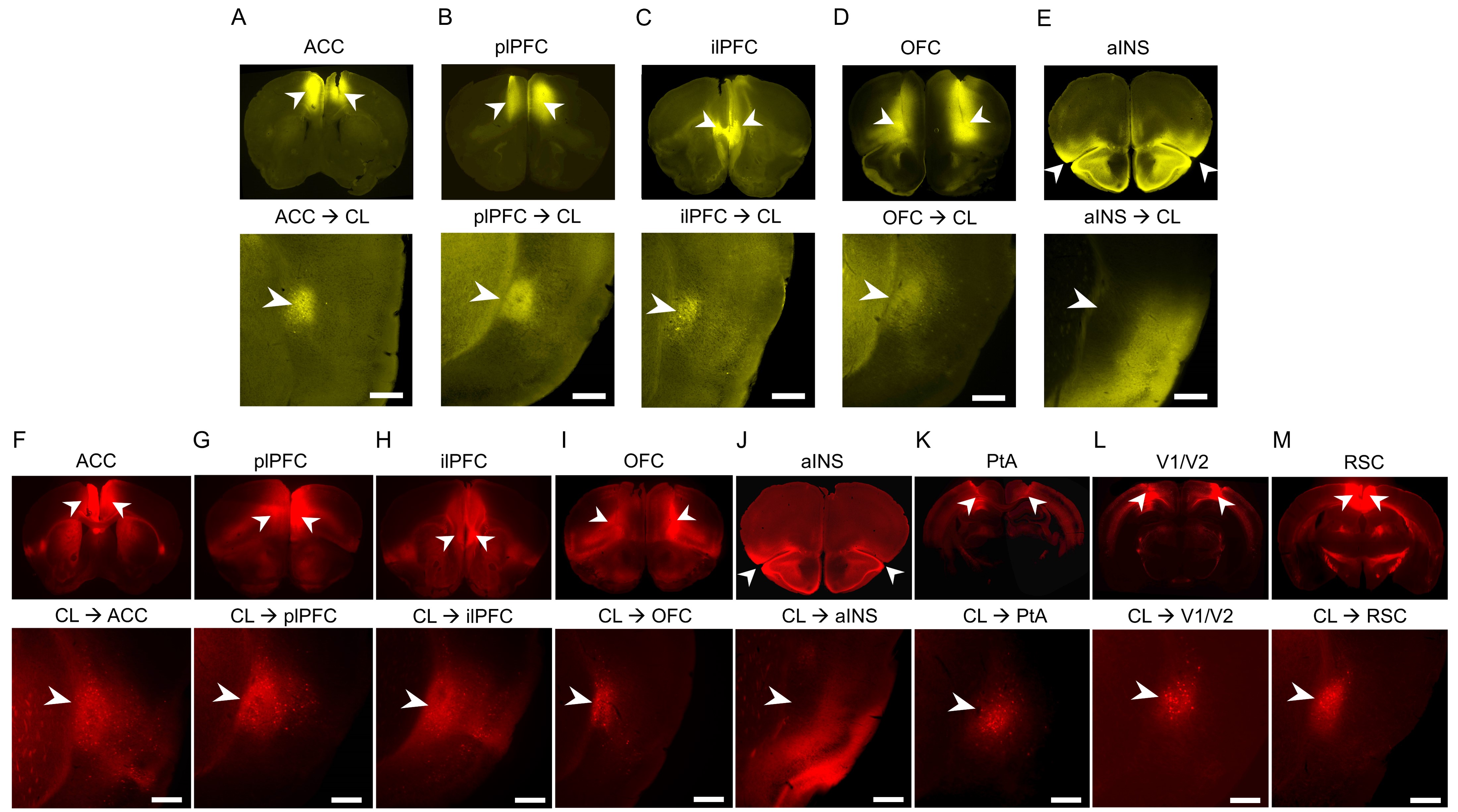

### Supplemental Figure S2

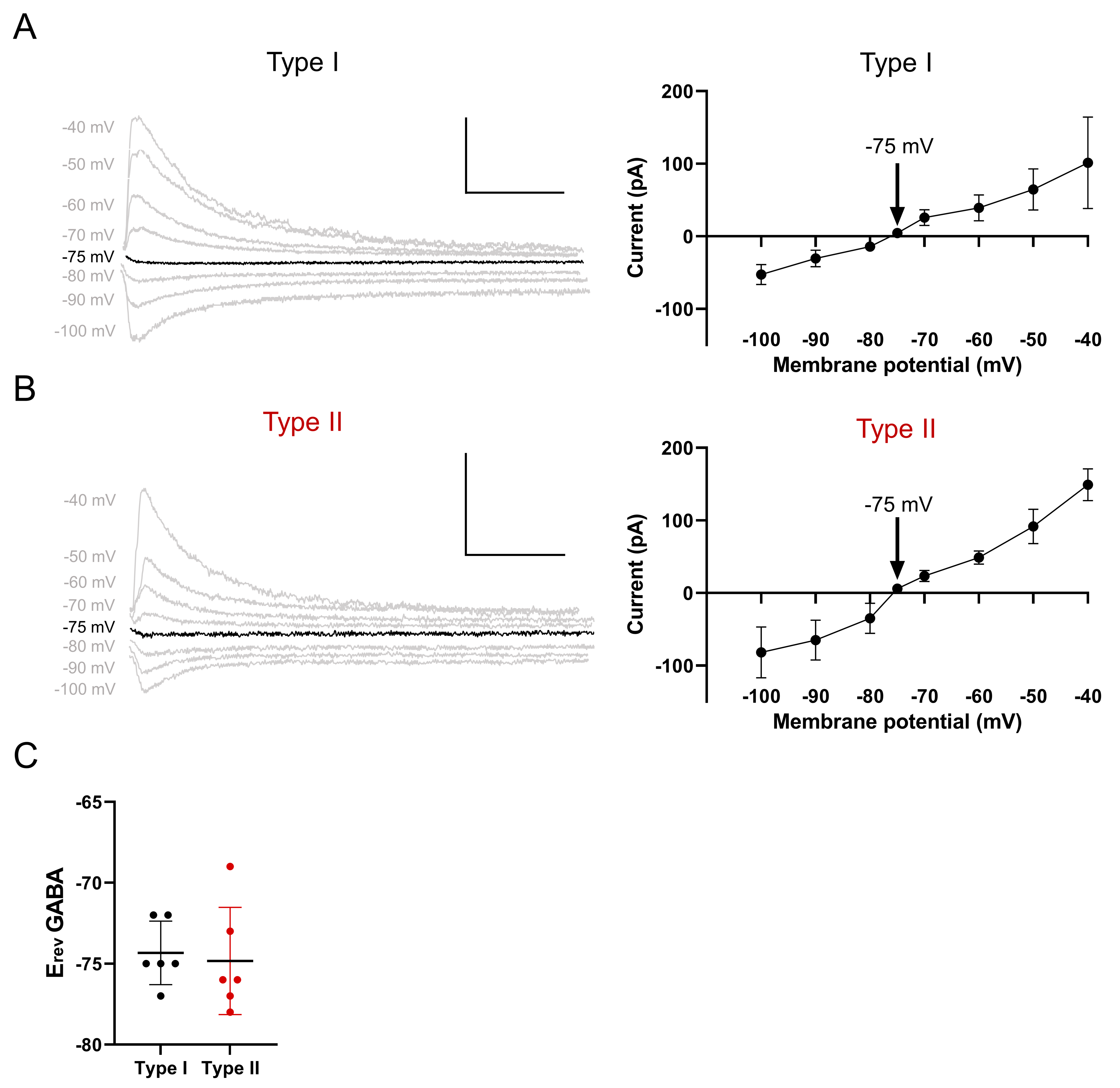

### Supplemental Figure S3

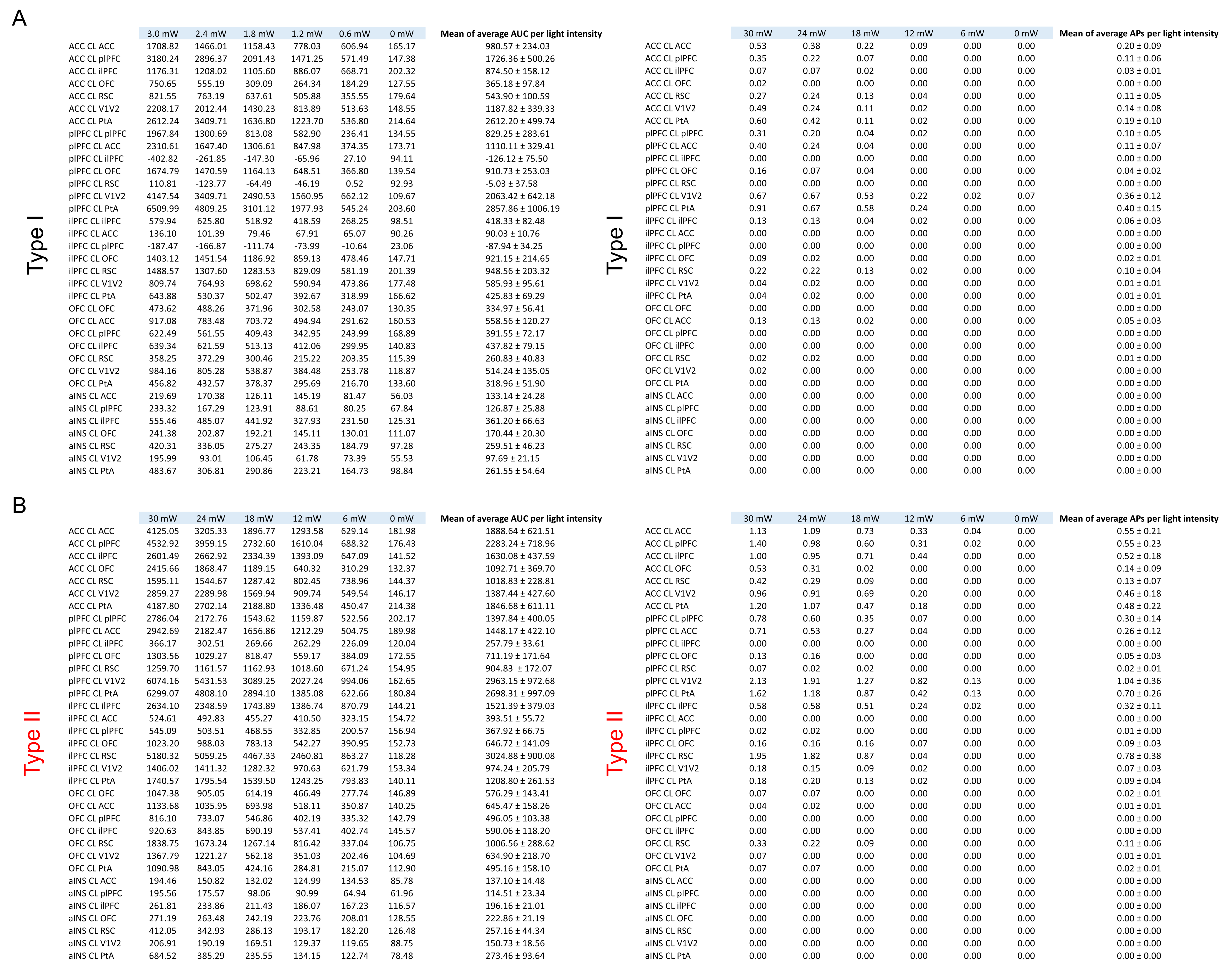

### Supplemental Figure S4

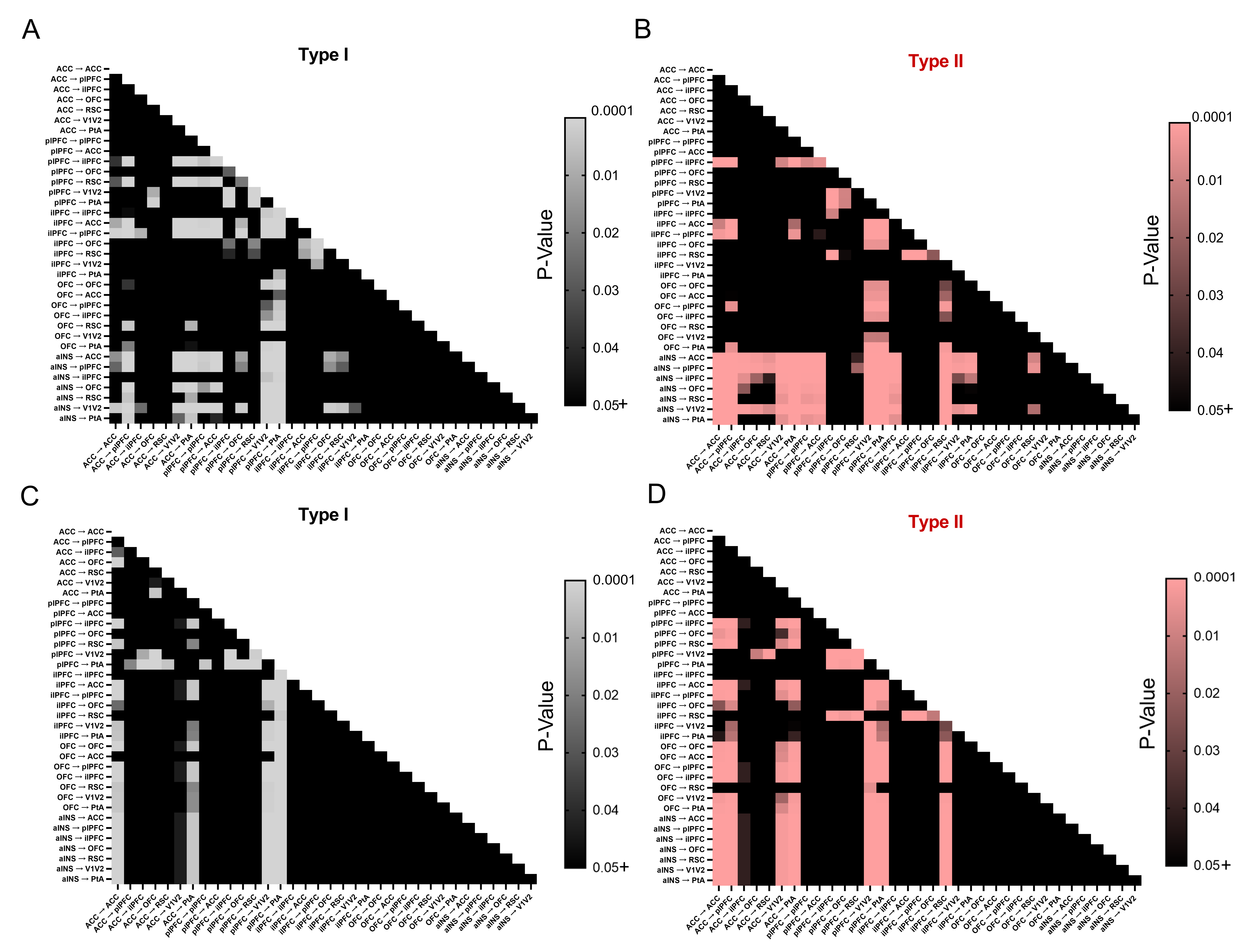
